# Within-subject Consistency of Paired Associative Stimulation as Assessed by Linear Mixed Models

**DOI:** 10.1101/434431

**Authors:** Myrthe Julia Ottenhoff, Lana Fani, Nicole Stephanie Erler, Jesminne Castricum, Imara Fedora Obdam, Thijs van der Vaart, Steven Aaron Kushner, Marie-Claire Yvette de Wit, Ype Elgersma, Joke H.M. Tulen

## Abstract

Paired associative stimulation (PAS) is a frequently used TMS paradigm that induces long-term potentiation in the human cortex. However, little is known about the within-subject consistency of PAS-induced effects. We determined PAS-induced effects and their consistency in healthy volunteers between two PAS sessions. Additionally, we assessed the benefit of applying linear mixed models (LMMs) to PAS data. Thirty-eight healthy volunteers underwent two identical PAS sessions with a >1 week interval. During each session, motor evoked potentials (MEPs) were assessed once before PAS induction and 3 times after at 30 min intervals. We did not detect any significant potentiation of MEP size after PAS induction. However, MEP size during PAS induction showed significant potentiation over time in both sessions (LR(1)=13.36, p<0.001). Nevertheless, there was poor within-subject consistency of PAS-induced effects both during (ICC=0.15) and after induction (ICC=0.04-0.09). Additionally, statistical model selection procedures demonstrate that a LMM with an unstructured covariance matrix better estimated PAS-induced effects than one with a conventional compound symmetry matrix (LR(34)=214.73, p<0.001). While our results are supportive of a high intra-individual variability of PAS-induced effects, the generalizability of our results is unclear, as we were only partially successful in replicating results from previous PAS studies typically showing potentiation of MEPs during and after PAS induction. We do, however, demonstrate that linear mixed models can improve the reliability of PAS-induced effects estimation.

## 1 Introduction

Synaptic plasticity is a fundamental process in our central nervous system, as it is essential for learning and memory (Caroni et al., 2012; Caroni et al., 2014). In addition, plasticity deficits are important in the etiology of many neurocognitive disorders (Klyubin et al., 2014; Srivastava and Schwartz, 2014). Synaptic plasticity is conventionally measured with invasive intraparenchymal electrophysiological techniques, which cannot readily be performed in human subjects. The development of transcranial magnetic stimulation (TMS) paradigms, such as paired associative stimulation (PAS) (Stefan et al., 2000), has enabled measuring plasticity-like effects in human subjects non-invasively, facilitating translation of findings from animal models to humans.

PAS is typically applied by pairing median nerve stimulation (MNS) with magnetic stimulation of the contralateral hand area of the primary motor cortex (M1). Consistent with the fundamental properties of spike-timing dependent plasticity (STDP) [7], when MNS precedes magnetic stimulations by 25ms, PAS stimulation induces a long-term increase in excitability of the M1 hand area, observed as an increase of motor-evoked potentials (MEPs) in the contralateral hand. In contrast, if the MNS precedes the magnetic stimulation by 10ms, the result is a long-term depression effect (Wolters et al., 2003). The resemblance to STDP is further strengthened by evidence that PAS-induced effects are dependent on the function of the *N*-methyl-D-aspartate (NMDA) receptor, known to be essential for long-term synaptic plasticity (Stefan et al., 2002).

Because of the similarity of PAS results to STDP experiments in rodents, PAS has emerged as a potentially very useful proxy for studying long-term synaptic plasticity in human subjects. However, PAS produces highly variable results between subjects (López-Alonso et al., 2014; Lahr et al., 2016; Wischnewski and Schutter, 2016), which is often attributed to the challenge of achieving similar levels of standardization as for animal experiments: environmental factors, lifestyle, experimental conditions and even genetic determinants have been suggested to influence the magnitude of the PAS-induced plasticity (Müller-Dahlhaus et al., 2008; Ridding and Ziemann, 2010; Wischnewski and Schutter, 2016). However, such factors only explain between-subject variability, whereas to our knowledge only one study examined the within-subject consistency (Fratello et al., 2006). More knowledge on this consistency is obviously important for studies that aim to follow human brain plasticity longitudinally.

Besides inter-and intra-individual variability, PAS studies show variable effect sizes between laboratories as well (Lahr et al., 2016; Wischnewski and Schutter, 2016). In addition to optimizing experimental procedures, some types of variability might be possible to account for by appropriate statistical modeling. PAS measurements generate relatively complex data, combining both repeated measures as well as a hierarchical data structure (i.e. multiple MEP size assessments per time point). In the last decades, linear mixed models (LMMs) have emerged as a statistical method that is specifically suited to handle such a data structure, reducing the chance of both false-positive and false-negative results (Aarts et al., 2014; Aarts et al., 2015). Additionally, LMMs are excellent for estimating reproducibility measures in the form of intra-class correlations. To date, however, LMMs remain to be sparingly applied to TMS data (Cash et al., 2015; Pedapati et al., 2015) and PAS-TMS data in particular (Cash et al., 2017).

In this study, we therefore assess the within-subject consistency of PAS-induced effects in healthy volunteers using two identical PAS sessions with an interval of at least 1 week, using LMMs.

## 2 Materials and Methods

### 2.1 Subjects

Thirty-eight out of 61 subjects were included in this study (reasons for exclusion are summarized in Table S1), who were recruited by advertising in the local community and on a Dutch research subject-recruitment website. Subjects were included if aged 18-40, right-handed according to the Edinburgh Handedness Inventory (Oldfield, 1971), in good health, medication free (excluding contraceptives) and able and willing to give written informed consent. Subjects were excluded if they were women lactating or pregnant, had a history of psychiatric illness and/or treatment, had a history of neurological illness or did not meet the international safety guidelines considering TMS (Rossi et al., 2009; Rossi et al., 2011). All subjects underwent the Wechsler Abbreviated Scale of Intelligence (WASI) (Wechsler, 1999) to determine their intelligence quotient (IQ) (Axelrod, 2002) for descriptive purposes. This study was approved by the Medical Ethical Review Board of the Erasmus MC Rotterdam, requiring study procedures to comply with the latest version of the Declaration of Helsinki.

### 2.2 Electromyography

Muscle activity was recorded from the left abductor pollicis brevis (ABP) muscle with electromyography (EMG), using Ag-AgCl electrodes in a belly-tendon montage. EMG signals were amplified using a universal amplifier (ANT Neuro, Enschede, The Netherlands) and digitalized at 5kHz for later offline analysis using Visor2 XT software (ANT Neuro, Enschede, The Netherlands). During measurements, a continuous EMG signal and trigger related EMG epochs were plotted at real time for online analysis, while applying a 50Hz notch filter and a 20-2000Hz bandpass filter.

### 2.3 Transcranial magnetic stimulation

Subjects were invited in the afternoon between 12 and 5.30 PM (Sale et al., 2007), were asked to not perform intense physical activities 24 hours prior to the measurement and to not smoke nicotine cigarettes or drink coffee on the day of the measurement. They were seated in a comfortable chair with their left arm resting on a pillow and were told to maximally relax their left hand during the measurement. Magnetic stimulations were applied using a figure-of-eight coil with an inner diameter of 27mm and outer diameter of 97mm, connected to a MagPro X100 with MagOption TMS device (MagVenture, Farum, Denmark). The coil was held tangentially to the left primary cortex and diverging 45° from midline. The electric field subsequently created in the cortex had a posterior to anterior direction.

To find the optimal position of the coil in order to maximally activate the ABP (the hotspot), TMS stimulations were randomly placed around a predefined reference point, defined as the location at 10% of the ear-to-ear span lateral to Cz over the right hemisphere. Data on coil location and position at every stimulation was collected using a neuronavigation system (ANT Neuro, Enschede, The Netherlands), allowing a precise definition of the angle and distance errors of every stimulation relative to the hotspot. All TMS procedures hereafter described are performed at the hotspot.

The resting motor threshold (RMT) was determined using a maximum-likelihood threshold hunting procedure (Awiszus, 2003). For this procedure, a MEP was defined as a signal with a peak-to-peak amplitude of ≥50μV. Subsequently, the stimulation intensity 1mV (SI1mV) was determined, which was the stimulation intensity of all subsequent stimulations. The SI1mV was defined as the percentage of maximal stimulation output (%MSO) of the TMS device that resulted in a mean MEP of 0.8 – 1.2 mV. For this purpose, trains of 10 magnetic stimulations at 0.1Hz at a chosen %MSO were performed until the criterion was met.

### 2.4 Paired associative stimulation

Subjects underwent two identical paired associative stimulation (PAS) sessions at >1 week apart. Baseline cortical excitability was assessed by applying a train of 20 magnetic stimulations at the SI1mV at 0.1Hz. Subsequently, PAS induction was performed by applying 200 paired stimulations of electric MNS preceding TMS by 25ms at 0.25Hz. After this plasticity induction phase, the cortical excitability measurement at baseline was repeated at three time points: immediately (Post 1), 30 minutes (Post 2), and 60 minutes (Post 3) (Figure 1A). MNS during the PAS-induction was applied at three times the sensory threshold using a bipolar bar electrode connected to a constant current stimulator (Digitimer Ltd., Letchworth Garden City, UK). If MNS surpassed the pain threshold, it was lowered to a painless but clearly noticeable level. The subject’s attention level was standardized by applying four randomly timed electric stimuli during PAS induction to the middle phalanx of the left thumb, and instructing participants upfront of PAS induction to focus their attention on their left thumb and report this number after PAS induction (Stefan et al., 2004). These stimulations were administered at two times the sensory threshold using a double ring electrode connected to a constant current stimulator (Micromed S.p.A, Mogliano Veneto, Italy).

**Figure 1.**
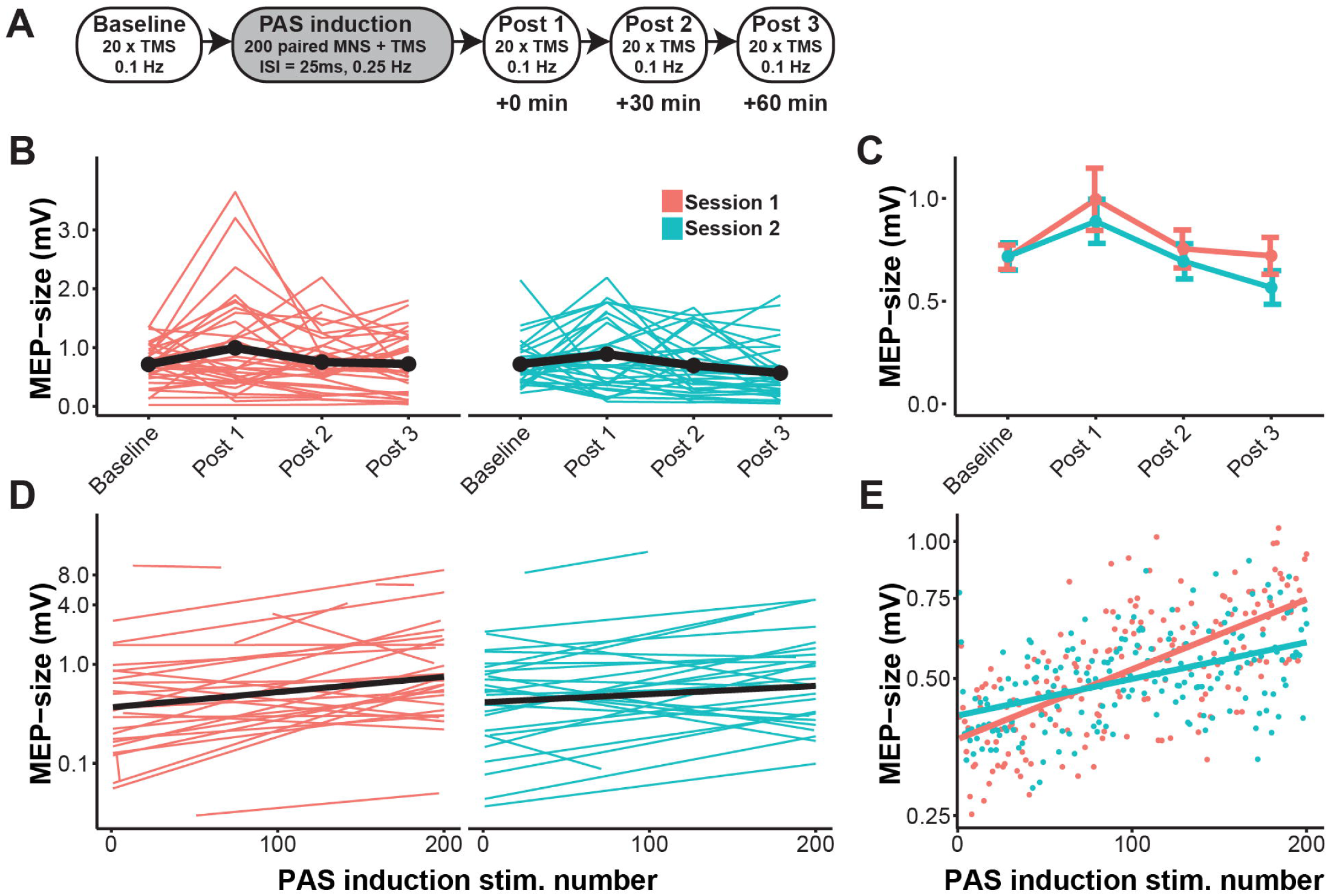
PAS-induced effects per session. Subjects underwent two identical PAS sessions spaced >1 week apart. (A) Schematic of one PAS session, in which the PAS induction is preceded by a baseline measurement consisting of 20 TMS stimulations, followed by a PAS induction phase consisting of 200 MNS-TMS paired stimulations, and 3 repeats of the baseline measurement at 30 min intervals. (B) The change in MEP size per session, where red line plots are individual medians of MEP size per time point in session 1 and blue line plots are those of session 2. The black line plots are means of individual median MEP size. (C) The change in MEP size over time for both sessions plotted together, where the connected dots represent means of individual medians and bars represent their standard error. Medians and means of individual medians were chosen as the best representative summary measure in B and C, as MEP size was not normally distributed in every data nest. (D) Linear regression lines through all MEPs during PAS induction per session (black lines) plotted over the linear regression lines through MEPs per individual (colored lines: red lines belong to session 1 and blue lines to session 2). (E) The change of MEP size over time during the PAS induction, with every dot representing the mean MEP size over all participants for that stimulation number. Lines are fitted linear regression lines per session. Note that in D and E the y-axis is log2-spaced.

### 2.5 Data analysis

The EMG signal for every magnetic stimulation applied was stored for offline analysis as epochs of −300ms to +300ms surrounding the TMS trigger. Using software programmed in LabVIEW (National Instruments, Austin, TX, US) pre-MEP noise, the maximal peak-to-peak amplitude and MEP onset were determined using a six-step data processing procedure:

1. Signals were linearly detrended.
2. The average amplitude value of the −300ms to −20ms before the TMS trigger was subtracted to create a zero-baseline.
3. To prevent ringing after filtering, the stimulation artefact was removed between −2ms to +4ms surrounding the TMS trigger, which was linearly interpolated. For PAS induction signals, the stimulation artefact of the MNS was removed similarly.
4. Filtering using both a 20-2000Hz bandpass filter and a 50Hz-notch filter.
5. Pre-stimulus noise quantification on a −25ms to +15ms time window surrounding the TMS trigger. After subtracting a 2^nd^-order polynomial fit, noise was defined as a peak-to-peak amplitude of >50μV or an SD of >15. Signals meeting these criteria were discarded for further statistical analysis.
6. MEP quantification, defined by the maximal peak-to-peak within a 20-48ms time window following the TMS trigger.

### 2.6 Statistical analysis

Statistical analyses were performed using R version 3.3.3 (R Development Core Team, 2018), supplemented with the nlme package (Pinheiro J, 2017). LMMs were used to estimate PAS-induced changes of MEP size, their correlations with baseline MEP size, and intraclass correlations (ICCs). For these LMMs, the dependent variable was MEP size, which was log2-transformed to better fit the assumption of normally distributed residuals. In addition, these LMMs were adjusted for log2-transformed angle and distance error.

We built Model 1 to estimate PAS-induced effects on MEP size *after induction* (Post 1, Post 2 and Post 3) within each session. This LMM included time point (categorical), session, and their interaction. The random effects included subject specific random effects for each time point in each session separately. An unstructured covariance matrix for the random effects was used (Model 1a) and was tested against the more restrictive compound symmetry structure (Model 1b).

Model 2 was built to estimate PAS-induced effects *during PAS induction*. This LMM included stimulus number (continuous), session and their interaction. Stimulus number was regarded as continuous time variable, as stimulations were equally spaced by 4 seconds in all PAS experiments. The model included subject specific random effects for stimulus number and session interaction and session. The eventual model was selected in three steps. First, we started out with a model using both natural cubic splines for stimulus number with three degrees of freedom and an unstructured covariance matrix (Model 2a). Second, to investigate the correlation structure, we tested Model 2a against a model with a compound symmetry structure (Model 2b). Last, to test whether the relation between MEP size and stimulus number was non-linear, Model 2a was tested against a model with a linear fit (Model 2c).

As a measure of within-subject consistency we calculated ICCs from LMMs that included session as an additional nesting level in the random effects. For the ICC of PAS-induced effects after induction, fixed effects and subject specific random effects of time point (categorical) were used (Model 3). For estimating the ICC of PAS-induced effects during PAS-induction over time, fixed effects as well as subject specific slopes for stimulus number (continuous time variable) were included (Model 4). Since the models used to calculate ICCs contained random effects for the respective time variables, the variation partition method was used (Goldstein et al., 2002). 95% confidence intervals (95%CIs) for each ICC were estimated using 500 bootstrap samples.

Likelihood-ratio tests were used to compare model fits and main effects of fixed effects. Descriptive statistics were performed using paired t-tests for normally distributed data, Wilcoxon Signed Rank tests for non-normal continuous data, a Chi-square test for categorical data or LMMs for data at the individual MEP level.

## 3 Results

### 3.1 Session characteristics

Thirty-eight individuals (22 women; median age 23, range 19-38; mean IQ 107±10SD) underwent two PAS sessions, which were spaced at least 1 week apart (median days between sessions was 14, IQR: 4). As displayed in Table 1, median starting time was significantly earlier in session 1 than in session 2, whereas both sessions did not differ in terms of baseline RMT, SI1mV, the level of attention during PAS induction or the angle and distance error relative to the hotspot.

**Table 1.**
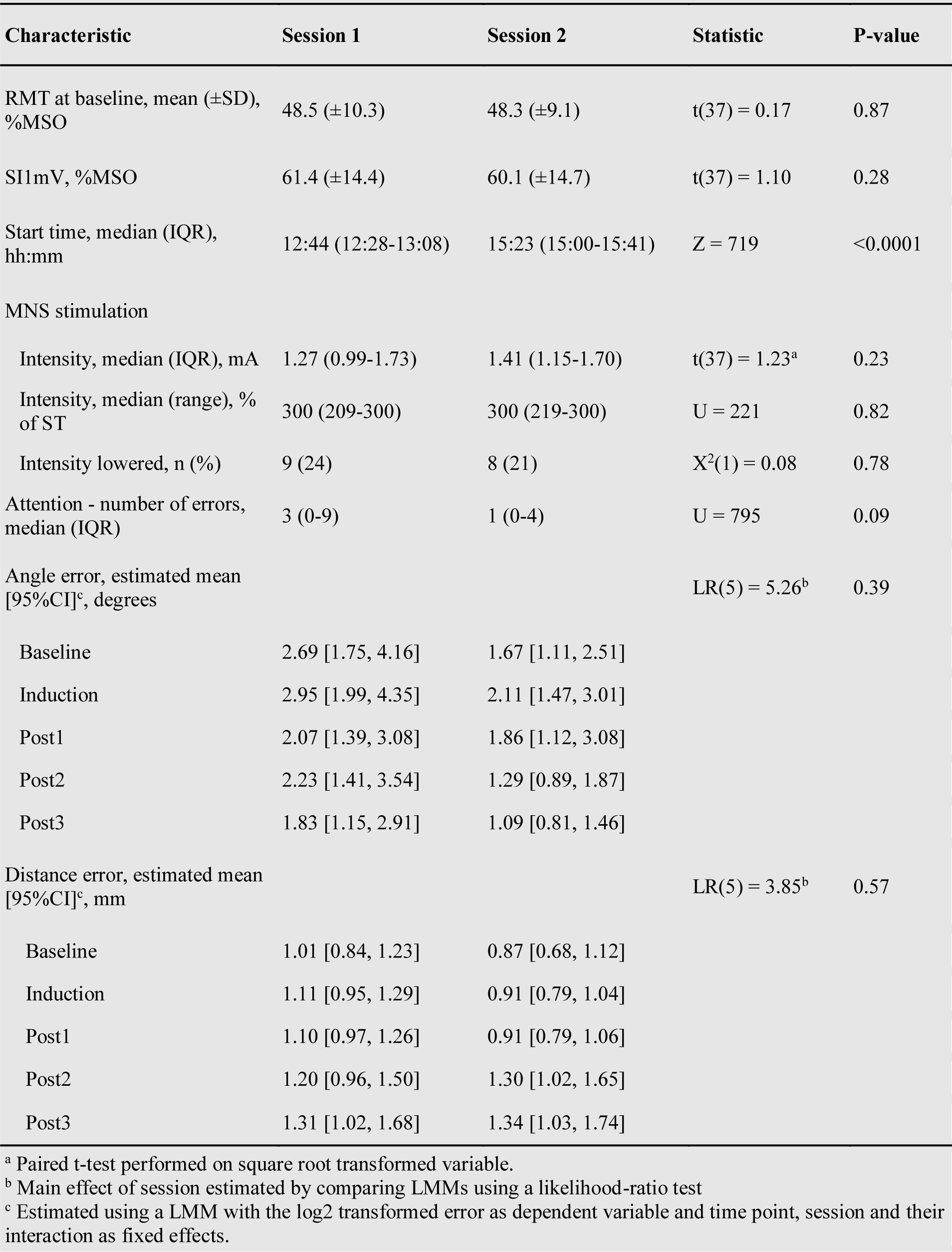
Session characteristics and comparisons.

To compare baseline MEP-size between session, we used the estimated means from Model 1a, 0.54mV (95%CI [0.43, 0.68]) for session 1 and 0.61mV (95%CI [0.53, 0.71]) for session 2, which were not significantly different (t(51650)=0.91, p=0.36). These model estimates are lower than expected, but it is important to note that the grand means are within the expected range: 0.91mV (±0.44 SD) for session 1 and 0.96mV (±0.36 SD) for session 2.

### 3.2 PAS-induced effects post induction

We determined the PAS-induced effect on MEP size at each post-induction measurement in each session. After filtering out MEPs with a noisy baseline, 5212 out of 6080 MEPs recorded (divided over 75 sessions and 38 subjects) could be used for this analysis. We estimated PAS-induced effects with a model with an unstructured covariance matrix that provided a superior fit to a model with a compound symmetry matrix (LR(34)=214.73, p<0.001). MEP size changed significantly over time (LR(6)=16.23; p=0.013), which was mainly driven by a negative effect on MEP size in Post 3 in session 2 (Table 2), instead of a positive effect on MEP size as is typically seen in PAS experiments. Additionally, individual trajectories of MEP size after induction were highly variable (Figure 1B). PAS-induced effects did not differ between sessions, as the interaction between time point and session was not significant (LR(3)=1.93; p=0.586), which is also reflected by the similar time courses in Figure 1C.

**Table 2.** Fixed effects of PAS induction on MEP size per post-induction time point and session estimated by linear mixed effect modelling.

| Variable | Session 1 |  |  |  | Session 2 |  |  |  |
| --- | --- | --- | --- | --- | --- | --- | --- | --- |
|  | B, % | 95%CI, % | t (5165) | P-value | B, % | 95%CI, % | t (5165) | P-value |
| All sessions |  |  |  |  |  |  |  |  |
| Post 1 | +16.83 | [-9.05, 50.07] | 1.22 | 0.22 | -3.99 | [-26.58, 25.55] | -0.30 | 0.77 |
| Post 2 | -10.00 | [-30.92, 17.24] | -0.78 | 0.43 | -23.92 | [-42.44, 0.55] | -1.92 | 0.05 |
| Post 3 | -14.84 | [-35.65, 12.69] | -1.12 | 0.26 | -31.65 | [-47.92, -10.30] | -2.74 | 0.006 |

The absence of significant PAS-induced potentiation is not consistent with most previous PAS reports (Wischnewski and Schutter, 2016). We, therefore, performed a subset analysis of sessions with a median baseline MEP size of ≥0.5 mV, as the observed low estimated baseline means could mean that the stimulation intensity during PAS induction was too low to induce robust potentiation. The ≥0.5mV subset contained 49 PAS sessions divided over 31 subjects (17 subjects retaining both sessions). Additionally, we explored a subset with <2 errors in the attention task, which contained 34 sessions divided over 28 subjects (5 subjects retaining both sessions), as subjects that had more errors could have poorer attention control leading to lower PAS-induced effects (Stefan et al., 2004). Both subsets showed similar PAS-induced effects compared to the full sample (Supplementary Figure S1 and Supplementary Table S2).

### 3.3 Potentiation during PAS induction

Next to the PAS-induced effects after induction, we determined the PAS-induced effect during induction. For this analysis, 9360 out of 15200 recorded MEPs were available due to filtering out MEPs with a noisy baseline, divided over 59 sessions within 34 subjects. Viewing the individual trajectories of MEP size development again indicates that there was high inter-individual variability (Figure 2C), which is reflected by the superior fit of the model with an unstructured covariance matrix to one with a compound symmetry covariance matrix (LR(8)=525.31, p < 0.001). The development of MEP size over time appeared to be linear (Figure 2D), supported by the fact that a model with a cubic fit was not superior to one with a linear fit (LR(4)=2.69, p=0.612).

**Figure 2.**
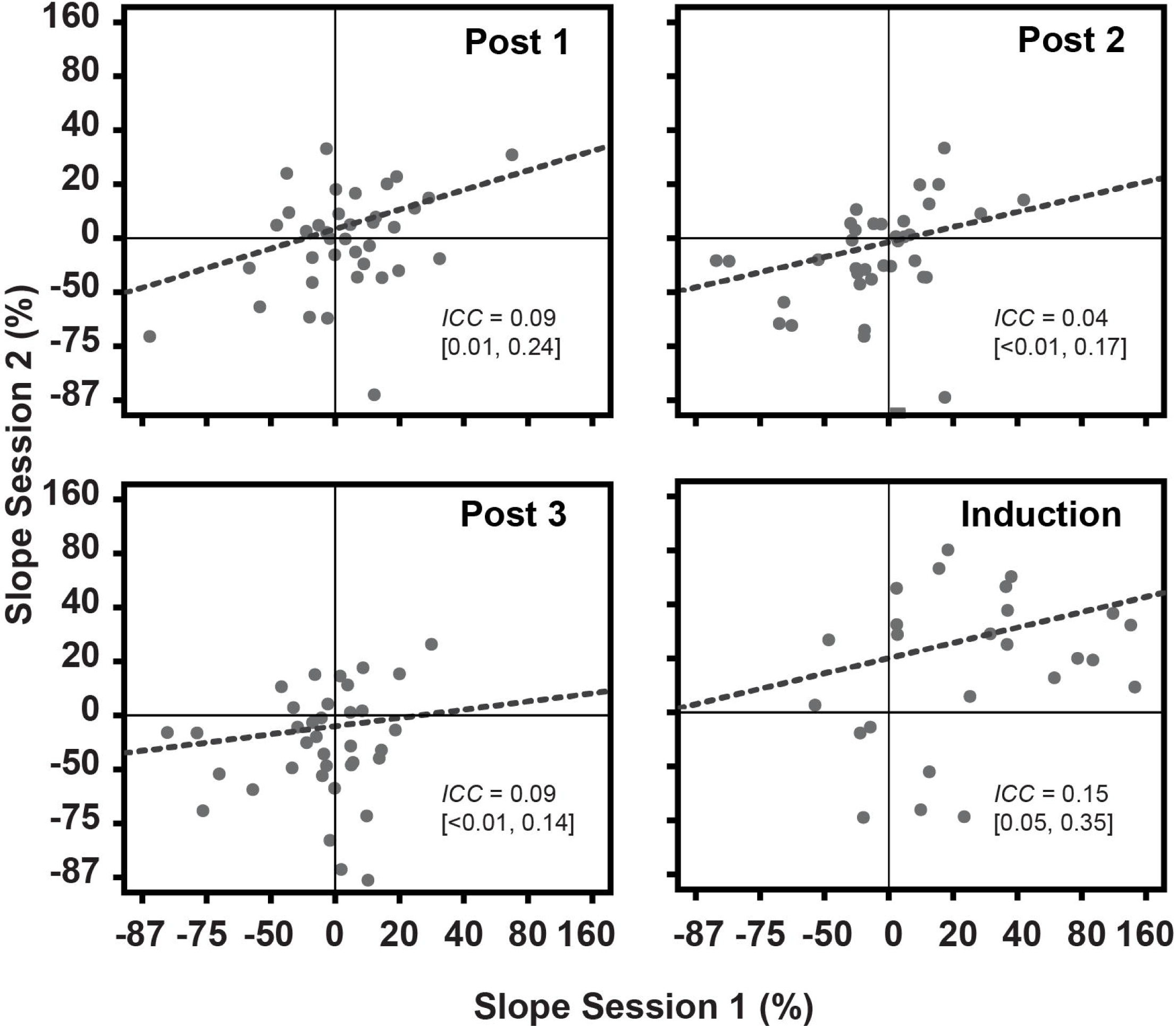
Intra-individual correlation of PAS-induced effects. Scatterplots of individual PAS-induced effects (grey dots) per measurement time point (Post 1-3 and Induction) of session 1 against those of session 2. These individual PAS-induced effects are the individual random slopes calculated by the models that were used to calculate the ICCs of PAS-induced effects reported in the Results section. Dashed lined represent the best linear fit. Note that the axes are log2-spaced.

Using the selected model with the unstructured covariance matrix and linear fit, we found that the estimated mean of MEP size at the start of PAS induction in session 1 (0.43 mV, 95%CI [0.27, 0.59]) did not differ from that in session 2 (0.44 mV, 95%CI [0.29, 0.66]) (LR(2)=0.967, p=0.617). There was a main effect of time (LR(1)=13.36, p<0.001), as a result of a significant positive increase of MEP size over time in both session 1 (+132%, 95%CI [+51%, +258%]) and session 2 (+79%, 95%CI [+19%, +169%]). However, there was no evidence of this time effect being different between sessions (LR(1)=0.87, p=0.35), reflected by the similar slope of the MEP size development in Figure 2D. There was a moderate negative correlation between MEP size at the start of PAS induction and the change in MEP size over time for session 1 (*r*=-0.51) and a weak negative correlation for session 2 (*r*=-0.41).

### 3.4 Consistency of PAS-induced effects

The within subject consistency of PAS-induced effects between the two sessions was poor: ICC^POST1^=0.09 (95%CI [0.01, 0.24]), ICC^POST2^=0.04 (95%CI [<0.01, 0.17]) and ICC^POST3^=0.04 (95%CI [<0.01, 0.14]) (Fig 3). Furthermore, the PAS-induced effects during induction showed a similarly poor within-subject consistency (ICC=0.15; 95%CI [0.05, 0.35]) (Fig 3), despite their significant potentiation at group level. The ICC of baseline MEP size before induction was poor (ICC=0.02; 95%CI [<0.01, 0.07]), as well as at the start of PAS induction (ICC=0.24; 95%CI [0.04, 0.42]). In contrast, the SI1mV did have a good within-subject consistency (ICC=0.88; 95%CI [0.83, 0.96]), as did the RMT at different time points (ICC^BASELINE^=0.85, 95%CI [0.77, 0.92]; ICC^POST1^=0.83, 95%CI [0.79, 0.90]; ICC^POST2^=0.85, 95%CI = [0.79, 0.92]; ICC^POST3^=0.85, 95%CI [0.78, 0.92]).

## 4 Discussion

We performed two identical PAS sessions in one group of healthy volunteers, resulting in pronounced potentiation over time during PAS induction, which was not consistent within subjects. PAS-effects after induction did not show the expected potentiation, and these effects were not consistent within subjects either. Additionally, we demonstrated that a linear mixed model with an unstructured covariance matrix provides the best model fit for our PAS data.

### 4.1 PAS-induced effects during and after induction

We found a significant increase of MEP size during PAS induction that shows striking resemblance to the increase in excitatory post synaptic potentials seen in STDP experiments in rodents (Froemke et al., 2010) and is consistent with previous human PAS studies (Dutra et al., 2016; Cash et al., 2017). From the animal studies, we know that the potentiation during plasticity induction correlates with the potentiation after induction. However, whether this increase in MEP size is a true proxy for NMDA-dependent LTP remains to be confirmed by sham-stimulation controlled studies and/or placebo-controlled NMDA-receptor antagonist intervention studies. It is noteworthy, however, that in our study MEP size at the start of PAS induction showed a negative correlation with PAS-induced effects during PAS induction. Namely, MNS during paired stimulations has a known acute inhibitory effect on MEP size, also known as short-latency afferent inhibition (Tokimura et al., 2000; Turco et al., 2018), lower MEP size at the start of induction could indicate more successful paired stimulations and, therefore, be related to a more prominent PAS-induced potentiation.

However, the significant potentiation during induction did not warrant significant potentiation after induction, which is not in line with most PAS studies (for review see (Wischnewski and Schutter, 2016)). This urged us to explored what factors could be responsible. First, our baseline MEP size appeared lower than the baseline in most PAS studies. It is, however, important to note that our grand means were within the expected range of MEP size and it is therefore unclear how our study compares to most PAS studies. Namely, many PAS studies solely report grand means without fully reporting whether both summarized and individual data are normally distributed. Nevertheless, due to this uncertainty, we have to consider that the low baselines observed here indicate that our stimulation intensity was possibly lower compared to most PAS studies, as several studies show that there is a positive correlation between this intensity and the PAS-induced effect (Meunier et al., 2012; Cash et al., 2017). Second, subjects that made more errors during the attention control task, could have had a negative effect on PAS-induced effects (Stefan et al., 2004). However, subsets of subjects with either a high baseline or few errors in the attention task did not show more PAS-induced potentiation, indicating that these factors are unlikely the cause of the absence of the potentiation of MEP size in our study.

Additionally, it is debatable whether our MNS was optimally performed, as some studies find a much stronger reduction of MEP size (Cash et al., 2015), while others suggest a reduction of similar degree (Elahi et al., 2012; Cash et al., 2017). This could be related to our use of a static 25ms MNS-TMS inter-stimulus interval opposed to adjusting this interval to the individual N20 peak timing (Ziemann et al., 2004). Another factor that could have contributed to the absence of PAS-induced potentiation is the known compromising effect of sleepiness on MEP size (Manganotti et al., 2004). As PAS is a lengthy experiment and subjects were not allowed to perform any type of physical activity or specific types of mental activity between post-induction time points, it is plausible that subjects became increasingly sleepy, masking potentiation effects. Unfortunately, although subjects were monitored to not fall asleep, we cannot support this speculation with actual measures of sleepiness, as there were not assessed.

### 4.2 Consistency of PAS-induced effects

The low ICCs found in this study seem to suggest that PAS-induced effects have a high intra-individual variability. One could, however, argue that the lack of significant post-induction potentiation compromises the validity of the consistency levels in this study. We did, however, show significant potentiation during induction, which showed similar low consistency consistent with (Fratello et al., 2006). They found equally poor intra-individual consistency of PAS-induced effects over two identical PAS sessions in a group of healthy volunteers (n=18), despite significant potentiation of post-induction MEPs at group level in each session. We, therefore, consider it not a given that the low ICCs are a consequence of the absence of a significant post-induction potentiation of MEP size.

Additionally, one could question whether our reported consistency would have been higher if we had eliminated MEPs classified as statistical outliers. As we took effort to eliminate MEPs based on confounding experimental conditions in the first place (pre-stimulus noise) and corrected for coil position errors, we regarded statistical outliers that remained in the dataset to be likely valid MEP measurements. Consequently, we view that retaining statistical outliers in our data set is important to reliably report ICCs.

### 4.3 Linear mixed models for PAS data

Our results provide insight in the potential advantage of LMMs for analyzing PAS data over conventional analysis methods. Most importantly, we show that using an unstructured covariance matrix provides a better model fit to our data than a compound symmetry matrix, for both estimating the PAS-induction effects during and after induction. This does not justify generalization of these findings for PAS data in general, but it does demonstrate that the estimation of PAS-effects benefits from the flexibility of the LMM in designing the covariance matrix. It is reasonable to suspect that PAS data with a complex multi-level and longitudinal structure inherits particular correlations between time points. Therefore, instead of ignoring the possibility of these correlations by using an analysis method that is restricted to the use of only the compound symmetry matrix (e.g. the RM-ANOVA), consistent implementation of LMMs for analyzing PAS data could improve reliability of results reported in PAS studies.

Another advantage of LMMs is that it does not require to summarize data per individual and time point (e.g. by averaging), but instead accounts for this data nesting by specifying random intercepts per nest. Data aggregation is problematic as it implies loss of information and, thus, statistical power. Additionally, if an incorrect data aggregation method is used, such as averaging while some data nests are left-skewed, PAS-induced effects could be overestimated.

### 4.4 Conclusion

While our results are supportive of a high intra-individual variability of PAS-induced effects, the generalizability of our results is unclear, as we were only partially successful in replicating results from previous PAS studies: we replicated the potentiation during the course of PAS induction, though this did not ensure significant potentiation after induction. Therefore, we cannot conclude to what extent PAS is a suitable outcome of human brain plasticity in longitudinal studies. It is worth emphasizing that our results demonstrate the benefit of linear mixed models for PAS data, as these models can reliably estimate PAS-induced effects despite the complex data structure and the various correlations between time points possible.

## Acknowledgements

We thank Dr. D. Rizopoulos for his helpful statistical advice, Dr. H.J. Boele for helping with developing the signal analysis software, Dr. H.G. van Steenis for technical support, and T. van Essen and D. Mani for their help preparing the data for signal analysis. This manuscript has been released as a Pre-Print at BioRxiv (Ottenhoff et al., 2018).

## Author Contributions

MO, LF, TV, SK, MW, YE and JT designed the study. MO, LF, IO and JC collected the data. MO and NE designed and performed the statistical analyses. MO drafted the manuscript. All authors interpreted the results, critically revised the manuscript for important intellectual content, and approved the final version of the manuscript.

## Data Availability Statement

The full dataset and linear mixed model syntax are available as Supplementary Data S1 (data set) and Supplementary Data S2 (syntax).

## References

Aarts, E., Dolan, C.V., Verhage, M., and van der Sluis, S. (2015). Multilevel analysis quantifies variation in the experimental effect while optimizing power and preventing false positives. BMC Neurosci 16(1), 94–94. doi: 10.1186/s12868-015-0228-5.

Aarts, E., Verhage, M., Veenvliet, J.V., Dolan, C.V., and van der Sluis, S. (2014). A solution to dependency: using multilevel analysis to accommodate nested data. Nature neuroscience 17(4), 491–496. doi: 10.1038/nn.3648.

Awiszus, F. (2003). TMS and threshold hunting. Suppl Clin Neurophysiol 56, 13–23.

Axelrod, B.N. (2002). Validity of the Wechsler abbreviated scale of intelligence and other very short forms of estimating intellectual functioning. Assessment 9(1), 17–23. doi: 10.1177/1073191102009001003.

Caroni, P., Chowdhury, A., and Lahr, M. (2014). Synapse rearrangements upon learning: from divergent-sparse connectivity to dedicated sub-circuits. Trends Neurosci 37(10), 604–614. doi: 10.1016/j.tins.2014.08.011.

Caroni, P., Donato, F., and Muller, D. (2012). Structural plasticity upon learning: regulation and functions. Nat Rev Neurosci 13(7), 478–490. doi: 10.1038/nrn3258.

Cash, R.F., Isayama, R., Gunraj, C.A., Ni, Z., and Chen, R. (2015). The influence of sensory afferent input on local motor cortical excitatory circuitry in humans. J Physiol 593(7), 1667–1684. doi: 10.1113/jphysiol.2014.286245.

Cash, R.F.H., Jegatheeswaran, G., Ni, Z., and Chen, R. (2017). Modulation of the Direction and Magnitude of Hebbian Plasticity in Human Motor Cortex by Stimulus Intensity and Concurrent Inhibition. Brain Stimul 10(1), 83–90. doi: 10.1016/j.brs.2016.08.007.

Dutra, T.G., Baltar, A., and Monte-Silva, K.K. (2016). Motor cortex excitability in attention-deficit hyperactivity disorder (ADHD): A systematic review and meta-analysis. Research in Developmental Disabilities 56, 1–9. doi: 10.1016/j.ridd.2016.01.022.

Elahi, B., Gunraj, C., and Chen, R. (2012). Short-interval intracortical inhibition blocks long-term potentiation induced by paired associative stimulation. J Neurophysiol 107(7), 1935–1941. doi: 10.1152/jn.00202.2011.

Fratello, F., Veniero, D., Curcio, G., Ferrara, M., Marzano, C., Moroni, F., et al. (2006). Modulation of corticospinal excitability by paired associative stimulation: reproducibility of effects and intraindividual reliability. Clinical neurophysiology: official journal of the International Federation of Clinical Neurophysiology 117(12), 2667–2674. doi: 10.1016/j.clinph.2006.07.315.

Froemke, R.C., Debanne, D., and Bi, G.-Q. (2010). Temporal modulation of spike-timing-dependent plasticity. Frontiers in synaptic neuroscience 2(June), 19-19. doi: 10.3389/fnsyn.2010.00019.

Goldstein, H., Browne, W., and Rasbash, J. (2002). Partitioning Variation in Multilevel Models. Understanding Statistics 1(4), 223–231. doi: 10.1207/S15328031US0104_02.

Klyubin, I., Ondrejcak, T., Hayes, J., Cullen, W.K., Mably, A.J., Walsh, D.M., et al. (2014). Neurotransmitter receptor and time dependence of the synaptic plasticity disrupting actions of Alzheimer’s disease Abeta in vivo. Philos Trans R Soc Lond B Biol Sci 369(1633), 20130147. doi: 10.1098/rstb.2013.0147.

Lahr, J., Passmann, S., List, J., Vach, W., Floel, A., and Kloppel, S. (2016). Effects of Different Analysis Strategies on Paired Associative Stimulation. A Pooled Data Analysis from Three Research Labs. PLoS One 11(5), e0154880. doi: 10.1371/journal.pone.0154880.

López-Alonso, V., Cheeran, B., Río-Rodríguez, D., and Fernández-Del-Olmo, M. (2014). Inter-individual variability in response to non-invasive brain stimulation paradigms. Brain Stimulation 7(3), 372–380. doi: 10.1016/j.brs.2014.02.004.

Manganotti, P., Fuggetta, G., and Fiaschi, A. (2004). Changes of motor cortical excitability in human subjects from wakefulness to early stages of sleep: A combined transcranial magnetic stimulation and electroencephalographic study. Neuroscience Letters 362(1), 31–34. doi: 10.1016/j.neulet.2004.01.081.

Meunier, S., Russmann, H., Shamim, E., Lamy, J.C., and Hallett, M. (2012). Plasticity of cortical inhibition in dystonia is impaired after motor learning and paired-associative stimulation. Eur J Neurosci 35(6), 975–986. doi: 10.1111/j.1460-9568.2012.08034.x.

Müller-Dahlhaus, J.F.M., Orekhov, Y., Liu, Y., and Ziemann, U. (2008). Interindividual variability and age-dependency of motor cortical plasticity induced by paired associative stimulation. Experimental brain research. Experimentelle Hirnforschung. Expérimentation cérébrale 187(3), 467–475. doi: 10.1007/s00221-008-1319-7.

Oldfield, R.C. (1971). The assessment and analysis of handedness: the Edinburgh inventory. Neuropsychologia 9(1), 97–113.

Ottenhoff, M.J., Fani, L., Erler, N.S., Castricum, J., Obdam, I.F., van der Vaart, T., et al. (2018). Within-subject consistency of paired associative stimulation as assessed by linear mixed models. bioRxiv, 434431. doi: 10.1101/434431.

Pedapati, E.V., Gilbert, D.L., Horn, P.S., Huddleston, D.a., Laue, C.S., Shahana, N., et al. (2015). Effect of 30 Hz theta burst transcranial magnetic stimulation on the primary motor cortex in children and adolescents. Frontiers in Human Neuroscience 9(February), 1–8. doi: 10.3389/fnhum.2015.00091.

Pinheiro J B.D., DebRoy S, Sarkar D and R Core Team (2017). “nlme: Linear and Nonlinear Mixed Effects Models”.).

R Development Core Team (2018). (Vienna, Austria: R Foundation for Statistical Computing).

Ridding, M.C., and Ziemann, U. (2010). Determinants of the induction of cortical plasticity by non-invasive brain stimulation in healthy subjects. The Journal of physiology 588(Pt 13), 2291–2304. doi: 10.1113/jphysiol.2010.190314.

Rossi, S., Hallett, M., Rossini, P.M., and Pascual-Leone, A. (2011). Screening questionnaire before TMS: an update. Clin Neurophysiol 122(8), 1686. doi: 10.1016/j.clinph.2010.12.037.

Rossi, S., Hallett, M., Rossini, P.M., Pascual-Leone, A., and Safety of, T.M.S.C.G. (2009). Safety, ethical considerations, and application guidelines for the use of transcranial magnetic stimulation in clinical practice and research. Clin Neurophysiol 120(12), 2008–2039. doi: 10.1016/j.clinph.2009.08.016.

Sale, M.V., Ridding, M.C., and Nordstrom, M.A. (2007). Factors influencing the magnitude and reproducibility of corticomotor excitability changes induced by paired associative stimulation. Exp Brain Res 181(4), 615–626. doi: 10.1007/s00221-007-0960-x.

Srivastava, A.K., and Schwartz, C.E. (2014). Intellectual disability and autism spectrum disorders: causal genes and molecular mechanisms. Neurosci Biobehav Rev 46 Pt 2, 161–174. doi: 10.1016/j.neubiorev.2014.02.015.

Stefan, K., Kunesch, E., Benecke, R., Cohen, L.G., and Classen, J. (2002). Mechanisms of enhancement of human motor cortex excitability induced by interventional paired associative stimulation. J Physiol 543(Pt 2), 699–708.

Stefan, K., Kunesch, E., Cohen, L.G., Benecke, R., and Classen, J. (2000). Induction of plasticity in the human motor cortex by paired associative stimulation. Brain 123 Pt 3, 572–584.

Stefan, K., Wycislo, M., and Classen, J. (2004). Modulation of associative human motor cortical plasticity by attention. Journal of neurophysiology 92(1), 66–72. doi: 10.1152/jn.00383.2003.

Tokimura, H., Di Lazzaro, V., Tokimura, Y., Oliviero, A., Profice, P., Insola, A., et al. (2000). Short latency inhibition of human hand motor cortex by somatosensory input from the hand. J Physiol 523 Pt 2, 503–513.

Turco, C.V., El-Sayes, J., Savoie, M.J., Fassett, H.J., Locke, M.B., and Nelson, A.J. (2018). Short-and long-latency afferent inhibition; uses, mechanisms and influencing factors. Brain Stimul 11(1), 59–74. doi: 10.1016/j.brs.2017.09.009.

Wechsler, D. (1999). “Wechsler Abbreviated Scale of Intelligence”. (San Antonio, TX: Psychological Corporation).

Wischnewski, M., and Schutter, D.J.L.G. (2016). Efficacy and time course of paired associative stimulation in cortical plasticity: Implications for neuropsychiatry. Clinical Neurophysiology 127(1), 732–739. doi: 10.1016/j.clinph.2015.04.072.

Wolters, A., Sandbrink, F., Schlottmann, A., Kunesch, E., Stefan, K., Cohen, L.G., et al. (2003). A temporally asymmetric Hebbian rule governing plasticity in the human motor cortex. J Neurophysiol 89(5), 2339–2345. doi: 10.1152/jn.00900.2002.

Ziemann, U., Ilic, T.V., Pauli, C., Meintzschel, F., and Ruge, D. (2004). Learning modifies subsequent induction of long-term potentiation-like and long-term depression-like plasticity in human motor cortex. J Neurosci 24(7), 1666–1672. doi: 10.1523/JNEUROSCI.5016-03.2004.

